# SEQPROC: an efficient, flexible, and concise tool for sequence geometry description and transformation

**DOI:** 10.64898/2026.07.28.741211

**Authors:** Noah Cape, Elan Fisher, Daniel Liu, Rob Patro

## Abstract

Complex sequencing protocols encode technical information in read structure and require accurate, efficient preprocessing. We introduce seqproc, which compiles concise sequence-geometry descriptions into execution graphs. Across four single-cell RNA-sequencing protocols, seqproc has the lowest mean runtime at every tested thread count and uses substantially less memory than the next-fastest tool. It has the highest F1 agreement with conservative structural references on all three discriminative chemistries and ties both alternatives on the 10x length-filter control. By separating protocol description from execution, seqproc makes complex read transformations compact, reusable, and efficient.

## 1 Background

Modern sequencing chemistries and protocols, particularly single-cell sequencing technologies, produce complex outputs that vary across platforms. Sufficiently complicated protocols could require dedicated processing pipelines after data generation in order to *decode* the technical information contained in the read [1–3]. For example, the sequencing reads may need to be processed in a specific way to extract the unique molecular identifiers (UMIs) and cell barcodes (CBs) associated with the sequenced fragment. These may be simple contiguous sub-sequences of a sequenced read, or may be spread across multiple distinct intervals denoted by the presence of specific motifs (linker or anchor sequences). These motifs themselves may be of variable lengths, and may undergo sequencing errors, requiring, for example, that they be matched inexactly.

The fact that different chemistries make use of different encodings of technical information has led to the creation of dedicated tools for pre-processing this data to identify and extract the relevant technical information in a way that can be easily specified and modified by the user [4–7]. While numerous sequence pre-processing tools have been created, we find them to either be (1) complex and verbose in their configuration, (2) limited in the technical pre-processing tasks they can accomplish, or (3) hindered by sub-optimal performance, or some combination thereof [8–11]. Some methods are simple to configure and specify, but can only process fixed length or exactly-matched sequences, while others can address a broader set of pre-processing tasks, but succumb to onerous configurations that tend to be verbose and complicated. Finally, since the pre-processing performed by these tools tends to be a computational kernel that is applied to *each fragment* (read) in a sequencing experiment, these kernels may be executed hundreds of millions or even billions of times when processing a single large sample. This brings performance considerations to the fore, and necessitates solutions that are not only flexible, easy-to-configure, and accurate in their processing, but also practically fast to execute.

In this paper, we introduce seqproc, a tool for pre-processing sequencing data. Our main contributions in seqproc comprise two main, and complementary, developments. First, we design and formally specify a descriptive, domain-specific language (DSL) called the Extended Fragment Geometry Description Language (EFGDL). It provides a concise yet powerful and flexible way to describe and manipulate both simple and complex fragment geometries, the ordered arrangement of functional elements (barcodes, UMIs, linker sequences, and cDNA inserts) within a sequenced read. Second, we introduce a novel, general, and efficient sequence matching and transformation library called antisequence that operates as the computational “engine” backing seqproc. antisequence provides a general framework for performing sequence matching and transformation operations through the modular composition of primitive operations in the form of a directed acyclic graph (DAG) of such sequence transformations. The result is a tool that allows for concise descriptions of fragment geometries and desired transformations that are compiled into an efficient computational pipeline that executes quickly (matching, and sometimes substantially exceeding that of existing state-of-the-art solutions). We demonstrate the utility of seqproc for pre-processing several different types of data, and compare it with existing tools for these tasks. Many tools have been introduced for performing technical pre-processing of sequencing data, with some tools designed specifically for certain data types and protocols [12–14], and others designed to allow more general and flexible formulation of the data being processed [8–10, 15]. Here, we focus primarily on tools that have been introduced with a focus on general and extensible pre-processing, and that have been primarily used for the pre-processing of high-throughput single-cell sequencing data.

splitcode [10] provides a command line tool and a graphical user interface to test configurations. The minimum necessary information to provide splitcode to process reads are the configuration file and the FASTQ file(s)^1^. In the configuration file, technical sequences are organized as tags and can then be extracted into named files. Flanking sequences can be extracted along with tags using the @extract keyword in the configuration file.

interstellar [9] provides a command line tool for sequence pre-processing. interstellar’s configuration file can be organized in logical blocks. This includes basic blocks like [general] which is used to specify a working directory and other blocks like [value-extraction] or [value-translation] where a read-structure is specified in accordance with the extended regular expression in the Python Regex library. Additionally, the [value-translation] block allows filtering of allocated sequence segments and defining the final read structure for outputted sequences. While the configuration of interstellar is simple and flexible after learning the necessary conventions, due to its core algorithms and implementation details, it runs much more slowly and requires much more memory than more recent alternatives.

The flexiplex [11] tool was introduced as a powerful read pre-processor focused mostly on the pre-processing and demultiplexing of long-read data (PacBio and ONT sequencing). It allows concise description of the manner in which read segments are meant to be matched and the fragments subsequently demultiplexed, using an extended regular expression-like language. While this descriptive language is concise and intuitive, it is somewhat limited in its generality for more complicated chemistries. Likewise, the blaze [16] tool has been introduced for barcode and UMI detection and pre-processing in ONT single-cell data (though it has also been successfully applied to PacBio data). It demultiplexes the reads into files with the UMIs and barcodes identified. It is designed for use with different 10x Genomics kits, and so the specification of the chemistry is very straightforward. However, this simplicity comes at the cost of generality, and so it does not readily support the description and processing of custom or complex chemistries.

read-structure [8] provides a Rust library for working with strings that describe how the bases in a sequencing run should be allocated into logical segments. It uses a “length” “specifier” design that is concise and simple. Although it is only a library for associating intervals to specifiers, and therefore cannot serve as a standalone pre-processing tool, the authors have also written software, like fqtk demux [17] which makes use of read-structures to parse and demultiplex sequencing reads. This tool does not output the FASTQ reads after parsing, and can only demultiplex reads against a list of known barcodes. Thus they fall outside the scope of comparisons we make here. Yet, our description language follows a similar philosophy of providing simple and concise inputs which, when the underlying geometry is itself simple, looks much like a read-structure. However, seqproc supports the specification of substantially more complex geometry and transformations, with the specification complexity only increasing when necessary to match the underlying geometry or processing being requested.

The recently introduced matchbox [15] tool provides a full DSL for sequence pre-processing, that is then interpreted and executed by the matchbox program. In terms of its execution model, matchbox is likely the closest in structure to seqproc. Yet, there are several key differences in both the design and implementation of these tools. Primarily, matchbox exposes an *imperative* DSL, where the user is responsible for describing *how* sequences are matched upon and transformed. On the other hand, seqproc is primarily a *descriptive* language, where the user describes the input geometry, along with appropriate labels and qualifiers for any intervals of interest, and then describes the desired output structure with details of precisely how the matching and transformation is done being left to the seqproc backend.

Beyond tools designed specifically for read pre-processing, several broader efforts have applied DSLs and specialized libraries to sequence analysis. SARVAVID [18] compiles a high-level DSL to efficient sequence analysis code through program synthesis techniques. The Seq language [19] extends Python with genomics-specific primitives and compiler optimizations for sequence processing pipelines. BioJulia [20] provides a composable library ecosystem for biological sequence manipulation within the Julia language. while these systems address sequence processing more broadly, seqproc targets a narrower niche, a declarative description and efficient pre-processing of read geometries as encountered in high-throughput sequencing experiments. This focus allows EFGDL to provide a more concise and purpose-built notation than general-purpose alternatives, while antisequence’s DAG-based execution model enables optimizations specific to the read pre-processing workload.

### 1.1 Contributions

We formally introduce seqproc in Section 5. Given sequencing data generated by a specific technology using a known protocol, seqproc allows one to describe the protocol in EFGDL, a descriptive language that we introduce here, which enables the construction of a high-performance pre-processing pipeline, executed by the antise-quence (also introduced here) engine. We show that seqproc is usually faster and substantially more memory-efficient than competing approaches. The separation of the EFGDL front end from the antisequence backend enables, simultaneously, a succinct description of the expected input and desired output and a highly-optimized engine to perform the requisite matching and transformation.

## 2 Results

In this section we analyze the efficiency and accuracy of seqproc by comparing the pre-processing of multiple widely used single-cell RNA sequencing (scRNA-seq) technologies [2, 3, 21]. We selected chemistries that range from structurally-simple pre-processing to those with complex geometries, requiring extraction and mapping of sequences within reads. Our benchmarks span four distinct data modalities, namely short-read paired-end (10x Chromium v2), short-read paired-end with complex anchored geometry (SPLiT-seq PE), long-read (LR-SPLiT-seq), and combinatorial indexing with variable-length barcodes (sci-RNA-seq3). Although all four are scRNA-seq protocols, the geometry and pre-processing requirements differ, from fixed-length barcodes and UMIs to anchored linkers and combinatorial variable-length barcodes. Furthermore EFGDL can be used to process read geometries for any protocol that can be described by EFGDL, and is not inherently limited to scRNA-seq data.

### 2.1 Experimental setup

All performance benchmarks were run on an otherwise-idle, dedicated lab node running Red Hat Enterprise Linux 9.8 (Linux 5.14), with one AMD EPYC 9555 processor (64 physical cores, 128 logical CPUs, x86_64), 377 GiB of RAM, and local NVMe-backed XFS scratch storage. Simultaneous multithreading was enabled on the host, but every run was pinned to 1, 4, 16, or 32 distinct physical cores and excluded their SMT siblings. The tested programs were seqproc 0.1.0 (commit cbc156b; antisequence commit f3b0f1e), matchbox 0.3.2, and splitcode 0.31.6. Each tool–dataset–thread combination was benchmarked three times on the full, uncompressed dataset, with tool order randomized within a frozen execution schedule. We report mean wall time with sample standard deviation and the largest maximum resident set size (peak RSS) observed across the three replicates. The Linux page cache was not flushed between runs; randomization prevents cache state from systematically favoring one tool. Timed runs included FASTQ reading, parsing, matching, filtering, and requested transformations, but directed requested sequence output to /dev/null, thereby excluding materialized-output writing and compression. Reference-agreement statistics were obtained separately from one 32-thread run per tool and dataset with FASTQ output materialized and structurally audited. Exact commands, configurations, input checksums, executable checksums, software revisions, and per-replicate measurements are available in the reproducibility repository.^2^

### 2.2 seqproc efficiently pre-processes scRNA-seq data

We compare seqproc to splitcode and matchbox for pre-processing data from four distinct single-cell protocols, representing a range of read lengths and library geometries. We analyzed SPLiT-seq paired-end (PE) short-read data from the original protocol validation [3] (SRR6750041), a PacBio HiFi long-read (LR) SPLiT-seq dataset [22] (SRR13948564), 10x Chromium v2 short-read data from PBMC samples [23] (SRR8315379), and sci-RNA-seq3 combinatorial indexing data [2] (SRR7827254). [24] maintains a library of all of the above, and many more, single-cell protocol geometries.

For each method, we either used the recommended configuration for the specific data or followed its documentation and translated the chemistry into a match-box or splitcode configuration.^3^ We aligned the workloads qualitatively. The 10x task is only a paired-read length filter: each tool rejects a pair when R1 is shorter than 26 bases and otherwise passes the pair through. The other three tasks both filter and project the biological components. The corrected SPLiT-seq PE layout is UMI(10)+BC3(8)+Linker 1(30)+BC2(8)+Linker 2(30)+BC1(8), and every tool projects the 34-base UMI+BC3+BC2+BC1 product. LR-SPLiT-seq analogously projects the same four components (32 bases in total; splitcode materializes them as component FASTQs) in either orientation, and sci-RNA-seq3 projects the natural 27– or 28-base BC1+BC2+UMI product without padding its variable-length BC1.

Exact matching semantics cannot be made identical without imposing tool-specific preprocessing. For PE and LR-SPLiT-seq, seqproc uses edit distance three for linkers and Hamming distance one against canonical barcode lists. splitcode uses substitution-only matching and, for LR data, requires separate forward and reverse-complement passes. matchbox does not provide substitution-only Hamming matching, and approximate adjacent captures can move biological field boundaries. For SPLiT-seq PE, we avoid this shift by matching canonical barcodes exactly, allowing edit distance three for the linkers, and adding explicit UMI/barcode length guards; a shifted linker hit cannot pass if it changes a captured field’s protocol length. For LR-SPLiT-seq, where an analogous guarded projection is not available, the primary boundary-safe configuration retains exact linker and canonical-barcode matching. Its declarative pattern searches an exact linker first, uses that hit to pin the adjacent fixed-length fields, and tests the captured terminal barcode against the canonical list; orientation-discriminating barcodes remain in the pattern. This anchor-first transcription avoids scanning every terminal-barcode candidate across each long read while retaining the same exact-match constraints. As alternative analyses, we report parenthetically in Tables 1 and 2, and explore further in the supplement, how matchbox performs with externally Hamming-expanded barcode lists. For the PE sensitivity, terminal BC1 remains an exact pattern anchor, while fixed-position BC3 and BC2 windows are tested for expanded-list membership after linker placement, avoiding an unnecessary three-list search. The LR sensitivity likewise uses the anchor-first plan, with expanded lists substituted for the canonical resources. Expansion substantially increases sensitivity, but still adds preprocessing outside the declarative configuration and greatly increases runtime. For sci-RNA-seq3, the matchbox configuration captures one variable-length prefix and guards it to the documented 9– or 10-base BC1 lengths. This avoids shifted output boundaries.

**Table 1.** Agreement with the conservative structural reference from one materialized-output, 32-thread run per tool and protocol, using the same executable versions and qualitatively aligned configurations as the performance campaign. Emitted (%) is the percentage of input reads emitted; Precision (%), Recall (%), and F1 are computed with respect to the per-protocol reference set *V*_total_ (Supplementary Section S2 and Supplementary Table S2). These quantities measure agreement with a declared structural reference, not biological ground-truth accuracy. *^†^* For SPLiT-seq PE, unparenthesized matchbox values use exact canonical-barcode matching, edit-distance-three linker matching, and explicit captured-component length guards; italic parenthetical values use the same linker budgets with externally generated Hamming-distance-one-expanded barcode lists. The expanded PE configuration retains terminal BC1 as an exact pattern anchor and checks the fixed-position BC3 and BC2 windows for list membership after linker placement. For LR-SPLiT-seq, both the primary and expanded-list configurations retain exact linker matching because an analogous boundary-safe guard was not available for the long-read geometry. The expanded LR sensitivity uses the same anchor-first plan as the primary configuration, substituting the expanded barcode resources for the canonical lists. All variants are evaluated against the same protocol-specific reference. *^♭^* On LR-SPLiT-seq, splitcode is evaluated as the set union of a forward and reverse-complement pass; duplicate reconciliation is performed analytically for this table but is not included in the timed result.

| Chemistry<br>(input size) | Method | Emitted (%) | Precision (%) | Recall (%) | F1 |
| --- | --- | --- | --- | --- | --- |
| SPLiT-seq PE<br>(77,621,181) | seqproc | 73.42 | 99.66 | 98.88 | **0.993** |
|  | matchbox<sup>†</sup> | 64.76 (*73.76*) | 99.78 (*99.58*) | 87.32 (*99.26*) | 0.931 (*0.994*) |
|  | splitcode | 68.92 | 99.96 | 93.10 | 0.964 |
| LR-SPLiT-seq<sup>b</sup><br>(5,764,421) | seqproc | 9.47 | 100.00 | 97.33 | **0.986** |
|  | matchbox<sup>†</sup> | 0.14 (*7.05*) | 100.00 (*100.00*) | 1.48 (*72.43*) | 0.029 (*0.840*) |
|  | splitcode | 7.44 | 99.76 | 76.31 | 0.865 |
| 10x Chromium v2<br>(234,382,218) | seqproc | 100 | 100 | 100 | 1.000 |
|  | matchbox | 100 | 100 | 100 | 1.000 |
|  | splitcode | 100 | 100 | 100 | 1.000 |
| sci-RNA-seq3<br>(22,088,821) | seqproc | 90.00 | 99.87 | 99.95 | **0.999** |
|  | matchbox | 88.55 | 99.87 | 98.34 | 0.991 |
|  | splitcode | 89.17 | 99.87 | 99.02 | 0.994 |

**Table 2.** Primary performance benchmark under qualitatively aligned workloads. Wall times are means *±* sample standard deviations over three randomized full-dataset runs; lower is better. Peak RSS is the largest replicate maximum resident set size at each method’s fastest tested thread count. Inputs were uncompressed and requested sequence outputs were directed to /dev/null. *^†^* For the two matchbox rows, italic parenthetical values are the wall time at 32 threads and peak RSS, respectively, for the Hamming-distance-one-expanded-barcode sensitivity configuration. The PE sensitivity uses the same fuzzy-linker budgets as its primary row, retains terminal BC1 as an exact anchor, and checks the fixed BC3 and BC2 windows for expanded-list membership after linker placement. The LR sensitivity retains exact linker matching and uses the same anchor-first plan as the primary LR configuration. Both parentheticals are single frozen /dev/null timings rather than three-replicate estimates. *^♭^* seqproc and matchbox search both long-read orientations natively. splitcode was timed as two sequential passes over the forward and precomputed reverse-complement inputs; the two pass times are summed, reverse-complement preparation is outside the timed interval, and no duplicate-reconciliation step is included.

| Chemistry<br>(input size) | Method | Mean wall time ± SD (s) | | | | Peak RSS at<br>fastest point (MiB) |
| --- | --- | --- | --- | --- | --- | --- |
|  |  | 1 thread | 4 threads | 16 threads | 32 threads |  |
| SPLiT-seq PE<br>(77,621,181) | seqproc | **116.487 ± 0.411** | **29.386 ± 0.029** | **25.611 ± 0.073** | **26.998 ± 0.173** | **50.0** |
| | matchbox<sup>†</sup> | 4,819.408 ± 69.523 | 1,273.112 ± 6.769 | 470.386 ± 2.289 | 358.173 ± 3.688 (*4,272.492*) | 11,085.5 (*12,665.1*) |
| | splitcode | 418.842 ± 4.026 | 116.283 ± 0.922 | 49.190 ± 0.192 | 48.716 ± 0.064 | 1,284.1 |
| LR-SPLiT-seq<sup>b</sup><br>(5,764,421) | seqproc | **30.520 ± 0.077** | **7.757 ± 0.030** | **3.519 ± 0.050** | **4.271 ± 0.051** | **58.0** |
| | matchbox<sup>†</sup> | 559.617 ± 2.268 | 196.187 ± 11.115 | 44.183 ± 2.453 | 25.442 ± 0.302 (*512.661*) | 67.0 (*347.1*) |
| | splitcode | 446.242 ± 6.575 | 114.265 ± 0.377 | 32.825 ± 1.099 | 17.239 ± 0.100 | 532.2 |
| 10x Chromium v2<br>(234,382,218) | seqproc | **102.928 ± 1.012** | **36.612 ± 0.087** | **42.314 ± 0.100** | **43.319 ± 0.218** | **14.0** |
| | matchbox | 1,075.199 ± 3.455 | 476.547 ± 3.052 | 473.005 ± 2.159 | 522.985 ± 5.658 | 40.5 |
| | splitcode | 142.211 ± 0.757 | 68.821 ± 0.236 | 68.847 ± 0.584 | 67.761 ± 0.380 | 1,250.8 |
| sci-RNA-seq3<br>(22,088,821) | seqproc | **16.118 ± 0.029** | **6.423 ± 0.150** | **6.456 ± 0.029** | **7.256 ± 0.838** | **15.0** |
| | matchbox | 233.997 ± 2.616 | 88.241 ± 0.958 | 72.204 ± 0.655 | 75.156 ± 0.834 | 3,113.0 |
| | splitcode | 26.966 ± 0.226 | 12.881 ± 0.000 | 11.664 ± 0.031 | 10.379 ± 0.050 | 1,833.7 |

To assess agreement of the emitted reads with independently specified protocol structure, we computed a conservative structural-reference set (*V*_total_) from the raw input of each protocol (Supplementary Section S2). Table 1 reports the percentage of input reads emitted, Precision (|*E* ∩ *V*_total_|*/*|*E*|), Recall (|*E* ∩ *V*_total_|*/*|*V*_total_|), and their harmonic mean F1. It is important to qualify that these are reference-agreement statistics, not biological ground-truth accuracy. Reads outside *V*_total_ remain unadjudicated. Performance is reported separately in Table 2, because the timing runs discarded sequence output and therefore are not themselves a source of recovery statistics. The protocol-specific references comprise 73.997% of SPLiT-seq PE pairs, 9.727% of LR-SPLiT-seq reads, 100% of 10x pairs, and 89.928% of sci-RNA-seq3 pairs. The same reference and denominator are applied to every tool within each protocol.

Each reference set is constructed from a separate, protocol-specific Python script rather than a benchmark competitor. It represents what might be expected in terms of precision and recall if one wrote a bespoke, protocol-specific Python script to perform parsing and extraction for each individual protocol to respect the desired approximate matching behavior while respecting the necessary structural constraints among the read components. It should not necessarily be interpreted as absolute “ground truth”, but represents a best-effort attempt at parsing each chemistry in the most reasonable and accurate manner, without respect to the computational (or development) cost of that preprocessing step.

seqproc has the lowest mean wall time on every primary dataset and at every tested thread count (Table 2). At each method’s own fastest tested setting, seqproc is 1.62× to 4.90× faster than the next-fastest tool, and at one thread the range is 1.38× to 14.62×. The best setting is workload-dependent. seqproc reaches its minimum at four threads for 10x Chromium v2 and sci-RNA-seq3, and at 16 threads for both SPLiT-seq workloads. At these per-method fastest settings, seqproc uses 14– 58 MiB peak RSS, while the next-fastest tool, splitcode, uses approximately 9–122 times as much memory. The expanded-list matchbox sensitivities demonstrate the computational cost of expressing approximate-whitelist acceptance through externally enumerated exact alternatives: their single-run, 32-thread wall times are 4,272.492 s for SPLiT-seq PE and 512.661 s for LR-SPLiT-seq, compared with 358.173 s and 25.442 s, respectively, for the primary canonical-list configurations. This distinction illustrates the value of native approximate-whitelist matching and optimized execution planning for declarative protocol descriptions.

For 10x Chromium v2, all three tools recover the identical complete input set, as all tools are capable of exactly filtering those input reads with technical read length ≥ 26. On sci-RNA-seq3, seqproc has reference-agreement F1 0.9991, compared with 0.9944 for splitcode and 0.9910 for matchbox, and all methods have high pairwise agreement, with a minimum Jaccard index of 0.9839 between read sets emitted by different methods. The reference excludes 25,960 reads with equal-best anchor placements at both permitted BC1 boundaries (i.e. considering a 9bp and 10bp barcode both yield an equivalently optimal parse); seqproc and splitcode retain all of these structurally ambiguous reads and matchbox retains 25,165, accounting for essentially all nominal false positives against the conservative *V*_total_.

On SPLiT-seq PE, *V*_total_ contains 57,437,503 of 77,621,181 pairs. seqproc recovers 98.88% (56,796,243 pairs) with an F1 score of 0.993. splitcode recovers 93.10% (53,472,550 pairs) with an F1 score of 0.964. Because matchbox does not currently support Hamming distance, its primary configuration matches barcodes exactly against the canonical lists. It permits edit distance three for both linkers and uses explicit captured-component length guards to prevent shifted approximate matches from changing UMI or barcode boundaries. This guarded configuration emits 50,266,141 pairs; 50,154,400 are in the reference, giving 99.78% precision, 87.32% recall, and F1 0.931. As a sensitivity analysis, we externally expanded each barcode list to include all Hamming-distance-one neighbors while retaining the same linker budgets and fixed component lengths. This emits 57,252,325 pairs with 99.58% precision, 99.26% recall, and F1 0.994, but its single 32-thread /dev/null timing was about 12× slower than the corresponding primary canonical-barcode mean and used more memory. The seqproc–splitcode emitted-set Jaccard index is 0.9298; for primary matchbox the Jaccard indices are 0.8702 with seqproc and 0.8225 with splitcode.

As an orthogonal comparison, we ran split-pipe v1.4.0, the vendor-provided preprocessing software, on all 77,621,181 SPLiT-seq PE input pairs using its native barcode-correction semantics and the protocol’s 94-nt barcode-read layout. split-pipe accepted 59,697,558 pairs (76.91%). seqproc emitted 56,991,381 pairs, of which 55,464,563 were in the vendor set, giving vendor-set precision 97.32%, recall 92.91%, F1 0.951, and Jaccard index 0.906. splitcode emitted 53,495,595 pairs, with an intersection of 52,904,336, giving vendor-set precision 98.89%, recall 88.62%, F1 0.935, Jaccard 0.878. The guarded fuzzy-linker matchbox configuration emitted 50,266,141 pairs, 49,549,557 of which were in the vendor set, giving vendor-set precision 98.57%, recall 83.00%, F1 0.901, and Jaccard index 0.820. Thus, seqproc has the highest F1 and Jaccard agreement with the full vendor set, while all three tools retain high vendor-set precision. Because split-pipe is licensed vendor software, its accepted-read set represents the preprocessing process for this data as best encoded by the technology’s vendor. While it is not a perfect stand-in for ground truth or biological accuracy, it represents a strong independent line of evidence about how these data should be preprocessed.

For LR-SPLiT-seq, *V*_total_ contains 560,699 of 5,764,421 reads (9.727% of the input) with a complete adjacent, structurally consistent sequence of matched elements (i.e. a cassette) in either orientation. seqproc emits 545,702 reads, all in the reference (100% precision), achieving an F1 score of 0.986. Dual-pass splitcode emits a 428,941-read set union, of which 427,895 are in the reference giving an F1 score of 0.865. The boundary-safe exact matchbox configuration emits 8,273 reads, all within the reference, but recalls only 1.48% (F1 0.029). This low recall is a consequence of the overall matching approach taken by matchbox. Native fuzzy adjacent captures can shift cassette boundaries, whereas exact matching requires every linker and barcode to be error-free. In addition, matchbox’s one-best policy selects an anchor placement before post-match whitelist guards are evaluated; a read containing both a valid cassette and an equally scoring decoy can therefore be rejected if the selected placement fails a guard. Allowing all equally good placements can emit duplicate records, so we retain the deterministic one-best policy. As a sensitivity analysis, we evaluated matchbox with Hamming-expanded barcode lists while retaining exact linkers and the anchor-first plan. This emits 406,132 reads, all in the reference, raising match-box recall to 72.43% and F1 to 0.840. The sensitivity still requires preprocessing of the barcode set by the user, includes variants with ambiguous canonical ownership, and remains substantially slower than native approximate-whitelist matching (see Table 2).

PacBio reads have random orientation, requiring both forward and reverse-complement searches. seqproc achieves this natively through #[match_ori(either)], and matchbox also searches both orientations in one run. splitcode does not support matching reads in both forward and reverse-complement orientation in a single pass. Therefore, its primary result is the set union of separate forward and reverse-complement passes. The timed two-pass result sums both passes but excludes preparation of the reverse-complement input and any duplicate-reconciliation step. The validity analysis unions read IDs to avoid double counting reads (716) accepted in both passes. Forward-only timing is presented as a controlled supplementary comparison (Supplementary Table S1).

### 2.3 Concordant downstream analyses results across reads processed by these tools

To test whether the preprocessing tools evaluated here under semantically matched configurations support the same biological conclusions, we processed all 77,621,181 SPLiT-seq PE read pairs (SRR6750041) with seqproc, splitcode, and the guarded fuzzy-linker matchbox configuration. The resulting sets contained 56,991,381, 53,495,595, and 50,266,141 read pairs, respectively. For every tool we placed the 66-nt cDNA sequence in read 1 and the 10-nt UMI followed by the three 8-nt barcodes in read 2, then quantified the products with STARsolo [25] 2.7.11b against mouse GRCm38 (Ensembl release 102) [26]. The primary common configuration uses the appropriate STARsolo CB_UMI_Complex geometry description with the EditDist_2 command line options for matching barcodes, which corrects the three barcode pieces independently. This is downstream correction, rather than a relaxation of the upstream read-retention rules. The upstream configurations already require each retained barcode to be exact or within Hamming distance one of a canonical barcode.

The three barcode-rank curves retain similar cell/empty transitions. Their inflections occur at ranks 258, 255, and 251 for seqproc, splitcode, and matchbox. The nonzero matrices contain 15,616, 14,838, and 14,361 barcodes, respectively (Supplementary Fig. S1). Pairwise per-gene Pearson correlation on log(1 + *x*) UMI totals ranges from 0.991 to 0.995, while per-barcode Pearson correlation ranges from 0.962 to 0.981 (Table 3). Thus, the guarded matchbox configuration produces aggregate and barcode-level quantification close to the other tools.

**Table 3.** Pairwise downstream concordance for the final SPLiT-seq PE outputs under the common STARsolo CB_UMI_Complex EditDist_2 configuration. The primary matchbox input uses canonical-barcode matching, edit-distance-three linker matching, and explicit captured-component length guards. Pearson correlations use total UMI counts on the log(1 + *x*) scale. Cell-type agreement, mean Jaccard over the six marker-based cell types, and adjusted Rand index (ARI) are calculated on the cells called by all three tools.

| Tool pair | Per-gene Pearson | Per-barcode Pearson | Cell-type agreement | Mean type Jaccard | Cluster ARI |
| --- | --- | --- | --- | --- | --- |
| SEQPROC/ SPLITCODE | 0.995 | 0.981 | 0.936 | 0.744 | 0.634 |
| SEQPROC/ MATCHBOX | 0.995 | 0.972 | 0.932 | 0.737 | 0.635 |
| SPLITCODE/ MATCHBOX | 0.991 | 0.962 | 0.936 | 0.732 | 0.606 |

At a threshold of at least 200 UMIs, seqproc, splitcode, and matchbox call 225, 220, and 221 cells; 220 cells are shared by all three tools. Cell-type composition is also stable (Figure 1A): neurons comprise 63.3–64.4% of cells and astrocytes 12.4–13.1%, with smaller oligodendrocyte, OPC, microglial, and endothelial populations. Among shared cells, 90.9% receive the same marker-based label from all three analyses. The pairwise results are similarly balanced (Figure 1B and Table 3): cell-type agreement is 0.936, 0.932, and 0.936; mean per-type Jaccard is 0.744, 0.737, and 0.732; and cluster ARI is 0.634, 0.635, and 0.606 for seqproc–splitcode, seqproc–matchbox, and splitcode–matchbox, respectively. Averaged over pairs, per-cell-type Jaccard is highest for neurons (0.968) and astrocytes (0.954) and lowest for the low-abundance OPC (0.436) and microglial (0.533) assignments; Supplementary Table S3 reports every pair rather than only these averages. These lower label-level values, alongside the stronger count correlations, show that low-signal discrete annotations are more sensitive to read-set differences than aggregate expression estimates.

**Fig. 1.**
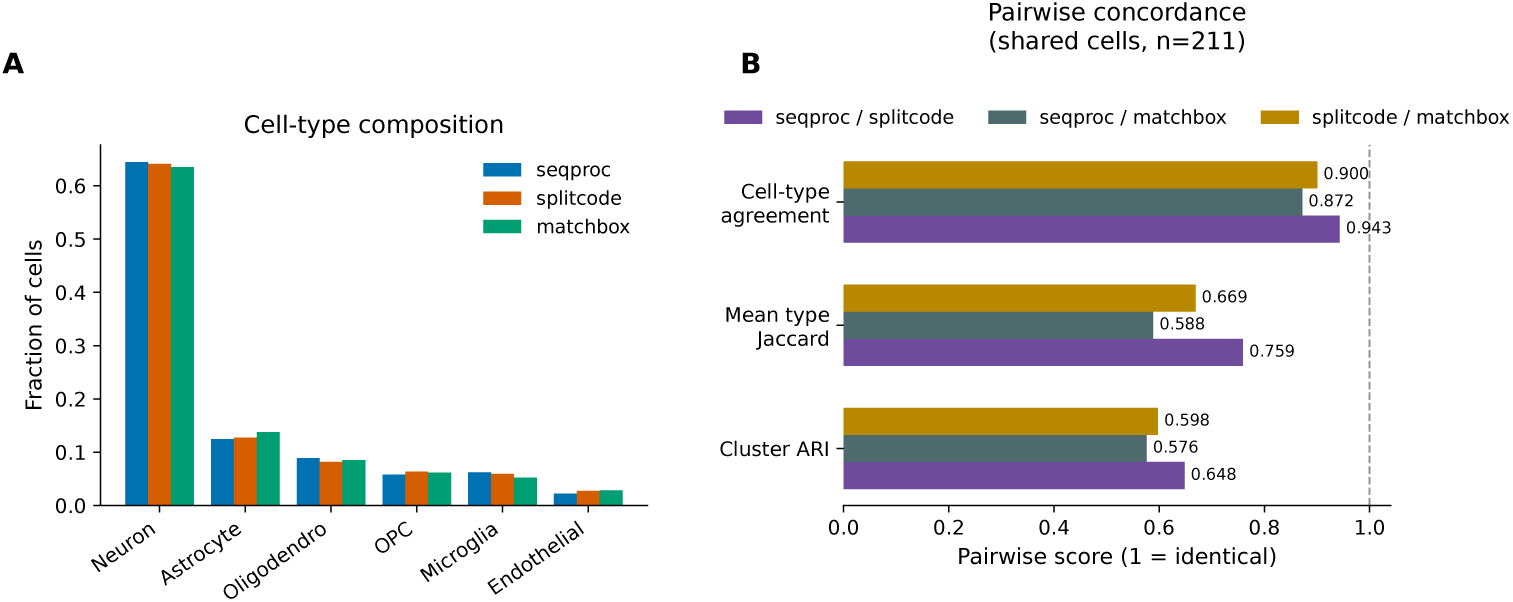
Downstream biological concordance for the final SPLiT-seq PE outputs, including the guarded fuzzy-linker matchbox configuration, quantified with STARsolo EditDist_2. **(A)** Marker-based cell-type composition among cells with at least 200 UMIs. **(B)** Pairwise concordance on the 220 cells called by all three tools. For seqproc–splitcode, seqproc–matchbox, and splitcode–matchbox, respectively, cell-type agreement is 0.936, 0.932, and 0.936; mean per-type Jaccard is 0.744, 0.737, and 0.732; and cluster ARI is 0.634, 0.635, and 0.606. The complementary all-tool cell-type agreement is 0.909, joint co-clustering agreement is 0.995, and tool mixing is 1.000.

**Fig. 2.**
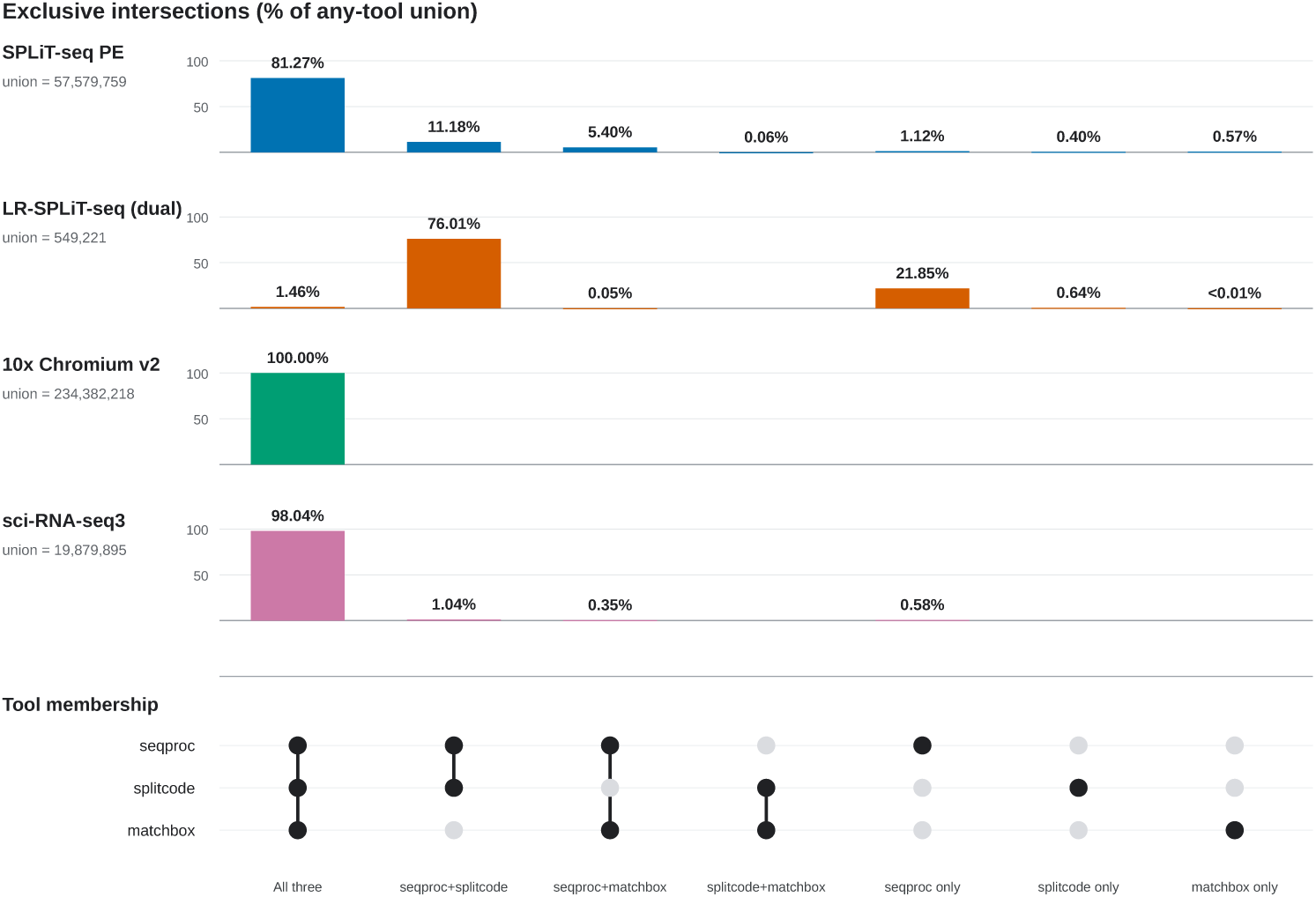
UpSet plot of emitted-read concordance from the same materialized 32-thread runs as Table 1. Within each dataset panel, bars show and label each mutually exclusive tool intersection as a percent-age of the union of reads emitted by any tool; each panel reports its union denominator. The matrix identifies which tools constitute each intersection. For LR-SPLiT-seq, splitcode is represented by the deduplicated set union of its forward and reverse-complement passes.

We also computed results processed with the 1MM STARsolo option to match barcodes from the whitelist and with the Hamming-expanded matchbox strategy. Those sensitivity analyses are reported in Supplementary Figs. S2–S4 and Supplementary Tables S4–S6. Across these alternatives, the main conclusion is qualitatively unchanged. Namely, gene-level quantification and major cell populations are robust, while barcode totals and rare marker-based labels reveal the expected sensitivity to barcode-correction and read-retention choices. On these data, seqproc also provides the strongest performance and memory profile (Table 2) while retaining the largest high-precision read set.

## 3 Discussion

The benchmarks show that a declarative read-geometry language need not trade performance for flexibility. Across protocols ranging from fixed-length barcodes to anchored, variable-length, and randomly oriented reads, seqproc has the lowest mean runtime at every tested thread count and a substantially smaller memory footprint than the next-fastest tool. Its high agreement with the conservative structural references and the concordance of downstream quantification indicate that these efficiency gains do not come at the expense of the evaluated outputs.

The comparison also has important limitations. The benchmark covers four single-cell RNA-sequencing protocols, and the protocol-informed reference sets are conservative structural references rather than biological ground truth. Reads outside those sets remain unadjudicated. The 10x task is intentionally a nondiscriminative length-filter control, and the degraded long-read library limits interpretation of absolute precision for every tool. In addition, matching semantics cannot be identical across programs without tool-specific preprocessing: the compared configurations differ in their support for edit distance, substitution-only matching, orientation handling, and adjacent approximate captures.

The downstream analyses are reassuring but should be interpreted at the appropriate level. Gene-level totals and major cell populations are stable across tools and sensitivity settings, whereas barcode totals and rare marker-based labels are more sensitive to correction geometry and read retention. The externally expanded matchbox barcode lists provide a useful sensitivity analysis, but their construction is preprocessing outside matchbox and shifts protocol knowledge into a generated auxiliary file. Quality scores are likewise not directly comparable because the tools differ in whether barcode qualities are observed or synthetic.

Several extensions could broaden the utility of seqproc. Additional built-in or user-defined annotations could support more context-dependent transformations. A community exchange of reusable EFGDL specifications could connect concise, machine-readable protocol descriptions with executable preprocessing pipelines and could draw on existing seqspec [27] descriptions. Further performance gains may be possible through broader runtime dispatch among existing matching implementations, operation fusion and reordering, dead-label elimination, and batching that adapts to graph cost, compression, and thread count.

## 4 Conclusions

seqproc provides a concise and flexible language for sequence geometry together with a high-performance execution engine for read preprocessing. Across the evaluated protocols it is the fastest tested tool at every thread count, uses substantially less memory than the alternatives, and has the strongest agreement with the conservative structural references on the discriminative chemistries. Its downstream results remain concordant with those obtained from competing tools. Separating the EFGDL description from the antisequence execution engine makes complex protocol specifications reusable while allowing backend optimizations to evolve with minimal disruption to users.

## 5 Methods

In this work we address the challenge of designing methods for the efficient parsing and transformation of potentially complex single-cell sequencing data. Here we formally describe the mathematical problem corresponding to the task, which our method, seqproc, aims to solve, and relate it to the existing theoretical models in the information parsing, extraction, and transformation literature, namely weighted finite-state transducers and spanners.

Let Σ be a finite alphabet. In the string-only setting (omitting quality scores and other record-level metadata) we model a sequencing read as a string *s* ∈ Σ*^∗^* and a dataset as a finite multiset or stream *U* of such strings. An interval of *s* is a half-open span [*i, j*), where 0 ≤ *i* ≤ *j* ≤ |*s*|; we write Int(*s*) for the set of all such intervals and *s*[*i, j*) for the corresponding substring.

A sequencing protocol specifies how a read is decomposed into named components, such as barcodes, UMIs, adapters, linkers, or biological sequence, and how those components are transformed. Let V be the finite set of component variables. A parse environment for *s* is a partial map *ρ*: V *⇀* Int(*s*), and we write Env(*s*) for the set of such environments.

**Definition** (Protocol description). A protocol description is a tuple P = (V, C, S*, E*)*, where* V *is a finite set of component variables,* C *is a finite set of structural constraints on their assigned intervals,* S *is a finite set of sequence-level constraints on the induced substrings, and E is an output expression evaluated on an input string together with a parse environment*.

The structural constraints in C specify how components are arranged, for instance their adjacency, precedence, fixed or bounded length, or position relative to a matched anchor. The sequence-level constraints in S specify the contents of intervals, including exact matching, bounded Hamming or edit distance matching, dictionary membership, or approximate dictionary membership. The expression *E* specifies the emitted output using transformations such as trimming, padding, removal, reversal, reverse-complementation, mapping, or filtering.

Given *s* and *ρ*, we write *s, ρ* |= *P* when *ρ* satisfies all constraints in C and S. The valid parses of *s* are

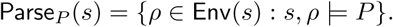

When approximate matching is used, parses may instead be cost-annotated,

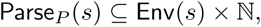

where (*ρ, c*) records a valid parse and its mismatch or edit cost. A protocol may also specify a priority, scoring, or tie-breaking rule that selects one parse.

For example, consider a protocol in which a read consists of an 8-nt cell barcode, a 12-nt UMI, an adapter sequence *a*, and a biological insert. Let V = {cb, umi, adapter, insert}. A valid parse *ρ* of a read *s* may satisfy *ρ*(cb) = [0, 8)*, ρ*(umi) = [8, 20), and, for some 20 ≤ *i* ≤ *j* ≤ |*s*|, *ρ*(adapter) = [*i, j*)*, ρ*(insert) = [*j,* |*s*|). The structural constraints require the fixed barcode and UMI positions and place the insert after the adapter, while the sequence-level constraint may require *d*_edit_(s[*ρ*(adapter)]*, a*) ≤ *k* for a fixed edit-distance threshold *k*. Thus, after an approximate adapter match is found, the remaining fields are interpreted relative to that anchor. One possible output expression is *E*(*s, ρ*) = *s*[*ρ*(cb)] · *s*[*ρ*(umi)] · revcomp^(^*s*[*ρ*(insert)]^)^, where · denotes concatenation. Thus, a protocol description is not merely a recognizer for valid reads, but a specification that induces interval assignments and transforms the associated substrings.

The recognized language of *P* is the set of strings admitting at least one valid parse *L*(*P*) = {*s* ∈ Σ*^∗^*: Parse*_P_*(*s*) ̸= ∅∥}.

**Problem 1** (Protocol Recognition). Given P and a finite multiset or stream U ⊆ Σ*^∗^, compute U_P_*= {*s* ∈ *U*: *s* ∈ *L*(*P*)}.

Problem 1 captures the case in which a protocol is used only to accept or reject reads. Sequencing pre-processing typically also requires recovering the intervals corresponding to protocol-defined components.

**Problem 2** (Protocol Parsing). *Given P and U, compute*

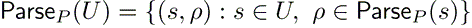

*or, in the cost-annotated case,*

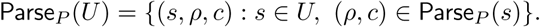

*If P specifies a priority, scoring, or tie-breaking rule, compute the selected parse for each read instead*.

Finally, let Δ be a finite output alphabet. The output expression *E* induces a partial transduction *T_P_*: Σ*^∗^ ⇀* Δ*^∗^* defined by *T_P_* (*s*) = Eval*_E_*(*s, ρ*) when *s* has a selected valid parse *ρ*, and undefined otherwise. If multiple parses are retained, then *T_P_*⊆ Σ*^∗^* × Δ*^∗^* is a relation.

**Problem 3** (Protocol Transduction). Given P with output expression E and a finite multiset or stream U ⊆ Σ*^∗^, compute*

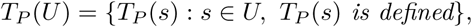

*or, in the relational case,*

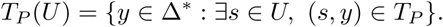

*Multiplicities and input order may be preserved when U is viewed as a multiset or stream*.

Problem 3 is the full string-level pre-processing task addressed by seqproc, which identifies protocol-consistent parses and emits the corresponding transformed strings or records. This formulation connects EFGDL to existing finite-state formalisms. Problem 1 is membership in the domain of a protocol. Problem 2 is closely related to regular spanners and variable-set automata, which map strings to assignments of variables to spans [28]. Bounded Hamming-distance matching, bounded edit-distance matching, and anchor-relative search can be modeled by finite-state or weighted finite-state automata or transducers when thresholds are fixed by the protocol [29, 30]. The output expressions in Problem 3 define transformations over captured intervals. Many of these are one-way finite-state transductions, while operations such as reversing or reverse-complementing an unbounded interval are more naturally captured by richer regular string-transduction models, such as two-way finite-state or streaming string transducers [31, 32]. Thus, the formal core of seqproc can be viewed as regular span extraction followed by regular string transformation, specialized to sequencing protocols. Efficient evaluation of regular spanners, including constant-delay enumeration after pre-processing [33], has also been studied.

To solve our particular instance of Problem 3, we introduce seqproc, which itself relies on EFGDL and antisequence.

### 5.1 seqproc

seqproc takes sequencing reads and an EFGDL specification (described in further detail below) as input. As an EFGDL to antisequence compiler, it compiles the specification into an antisequence program and streams the input reads through that program to yield the processed reads. Figure 3 outlines a seqproc workflow. The key innovation of seqproc is modularizing the solution of Problem 3 in our context, relegating the specification of *what* to match and transform to EFGDL and *how* to match and transform to antisequence.

**Fig. 3.**
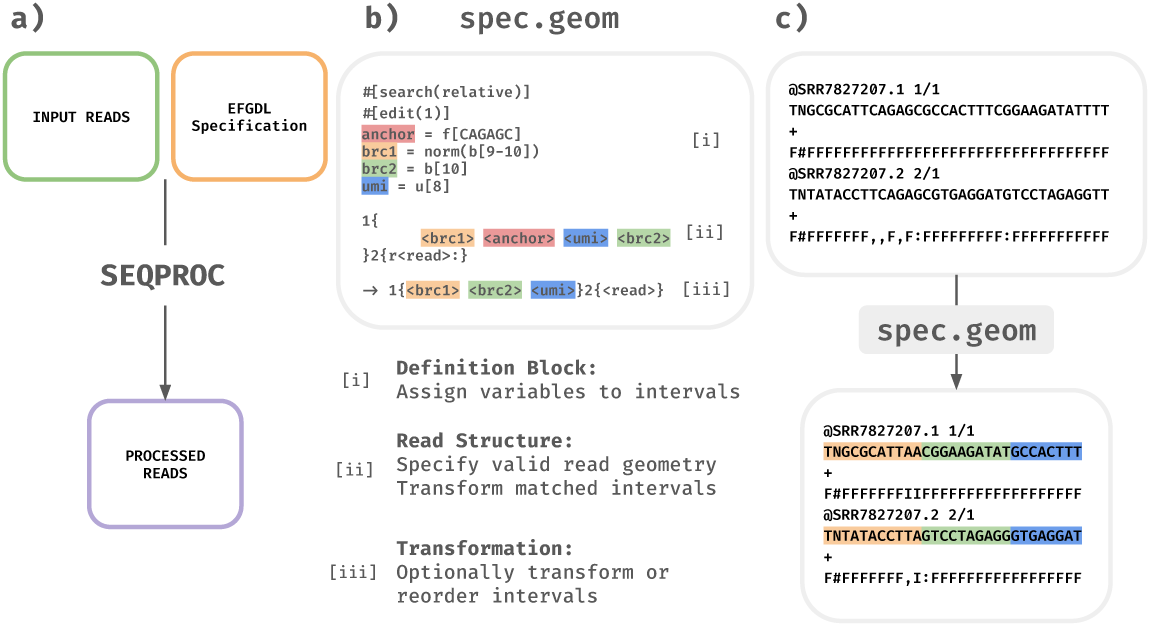
**a**) A cartoon example of how one would use seqproc, giving it both input reads and an EFGDL specification by which the input reads will be matched and possibly transformed. seqproc then operates on the reads using an antisequence execution graph and outputs, in a streaming fashion, those reads which passed the filtering. **b)** An EFGDL idiomatic specification for parsing and transforming sci-RNA-seq3 data [2] where intervals to be matched are named in the definitions, each interval is specified in the read structure and finally the extra information is removed and matched intervals are re-ordered for post-processing. **c)** An example of how some reads would be processed based on the EFGDL specification.

### 5.2 EFGDL

The Extended Fragment Geometry Description Language (EFGDL) is a descriptive language for composing the fundamental searching, matching, capturing and transforming operations for solving Problem 3. From the perspective of single-cell sequencing protocols, it is a language for describing the specific layout of information such as barcodes and unique molecular identifiers (UMIs) in a read as well as for describing how to optionally transform these intervals after locating them. seqproc parses the EFGDL specification using the Chumsky [34] parser combinator library. Chumsky uses parser combinators to generate recursive descent parsers that are capable of processing parsing expression grammars (PEGs) [35].

An EFGDL specification has three distinct components. *a*) **definitions**: where intervals are defined and associated with a label which can be referred to later. *b*) **read structure**: where intervals are chained together to define the grammar against which reads are to be matched. *c*) **transformation**: where intervals can be removed, reordered and otherwise transformed after being matched with labels. Only the presence of a read structure is necessary for a valid EFGDL specification. Definitions make read structures easier to understand, and transformations provide a concise syntax for removal, reordering or modification of labeled intervals. EFGDL has an extensive set of primitive transformations (Supplementary Table S7) that can be composed with specified intervals anywhere in an EFGDL program, described in more detail later.

EFGDL is a primarily descriptive language designed with two key goals, for the language to *1*) specify *what* to parse and transform in a sequence, and *2*) give a simple format to describe a sequencing protocol. For these reasons, the idiomatic way to write an EFGDL specification would be to declare all relevant intervals with descriptive names as definitions, then in the read structure refer to the defined intervals, followed by an optional transformation if intervals should be removed or re-ordered or modified. See the EFGDL specification in Figure 3 for an example of an idiomatic specification of the sci-RNA-seq3 protocol [2].

EFGDL intervals are the building blocks that specify which sequences are accepted by seqproc and which are filtered out. There are four types of EFGDL intervals *1*) **fixed sequence intervals** *2*) **fixed length intervals** *3*) **variable length intervals** and *4*) **unbounded length intervals**. To further describe or transform an interval, there are six interval specifiers, exactly one of which must be used to specify any declared EFGDL interval. These are **barcode** (b), **sample barcode** (s), **UMI** (u), **discard** (x), **read** (r), and **fragment** (f), where the specifier’s key-letter is written in parentheses to denote that interval type in an EFGDL specification. The fragment specifier is reserved for the fixed sequence interval and vice versa. All other specifiers and intervals can be combined in an EFGDL specification if desired. If an interval specified with a discard specifier is matched, that interval is removed from the read (provided the read itself is not filtered out). The sample barcode specifier (s) labels sample-of-origin barcodes in multiplexed protocols such as 10x Genomics Flex, where a probe barcode identifies the sample of origin and the cell barcode identifies the cell of origin, so that an identical cell barcode appearing in two different samples denotes two distinct cells. When used idiomatically, specifiers enhance the readability of the protocol and simplify the description of more complex transformations.

Writing valid EFGDL specifications requires using intervals to build an unambiguous grammar. Any variable or unbounded length interval must be anchored to the right by a known length set of intervals ending in the termination of the read or ending in a fixed sequence interval. The reason for this is that seqproc greedily matches intervals positionally, scanning left to right and binding each variable or unbounded length interval at the first (leftmost, by default) position of its right anchor; without such an anchor the boundary is undetermined. Optionally, users can apply annotations that change this behavior, as outlined below. When seqproc encounters a variable or unbounded length interval, it looks to the intervals on its right to determine how many bases to assign to it. For details on how seqproc determines and handles potential ambiguity, see Supplementary Section S1.

EFGDL’s interval transformations let seqproc pre-process both simple and complex geometries, emitting them in a concise, uniform structure suited to downstream analysis. Like a statically typed programming language, EFGDL transformations can only transform specific types of intervals. Because Chumsky supports error recovery, seqproc validates transformation arguments and reports any type mismatches before processing any reads. For example, a discard specified interval cannot be an argument to any transformation, because once it is matched, its subsequence is slated for removal from the output and is therefore unavailable to transformations. Furthermore, transformations can be *composed* to form more complex transformations.

Given an interval as an argument, each transformation (Supplementary Table S7) has one of two functions; it either modifies the interval’s associated subsequence, or modifies how the interval is eventually associated with a subsequence. Transformations like pad and rev are applied after a subsequence has already been associated with its respective interval argument, while matching modifiers control how an interval is matched to a subsequence and are specified using annotation syntax.

Annotations (with syntax inspired by the Rust programming language [36]) are placed before an interval definition using #[name(args)] notation. The annotation #[hamming(n)] or #[edit(n)] controls the matching tolerance for a fragment specified fixed sequence interval. #[search(relative)] modifies how seqproc locates a fragment within a read, searching for the anchor anywhere in the read rather than binding it at the leftmost feasible position. Annotations also compose. For example, lines 5-7 in Figure 4 define a linker that is located via relative search with edit distance tolerance 3, where #[search(relative)] controls positioning and #[edit(3)] controls matching tolerance. The annotation system is designed for extensibility, so that new matching strategies or metadata capture mechanisms can be introduced by adding new annotation types without modifying the core EFGDL grammar or existing specifications. This separation of matching behavior from interval definition ensures that EFGDL specifications remain forward-compatible as new sequencing technologies and pre-processing requirements emerge. Supplementary Table S7 gives an exhaustive list of the transformations currently available to a user writing an EFGDL specification. Property-like annotations use assignment syntax. In particular, equal-best matches against multiple distinct whitelist or mapping entries can be controlled with #[ambig_policy = value]. Exact duplicate list rows are normalized before matching and conflicting duplicate mapping rows are rejected, so this policy concerns genuine equal-best matches rather than duplicate input lines. Filters accept equal-best set membership by default, whereas mapping operations conservatively use their no-match fallback. Users can instead select accept, input-ordered first, no_match, deterministic random(seed = N), quality-aware quality(min_delta = N), or error. The quality-aware policy compares summed Phred scores at mismatch positions and accepts the best candidate only when it exceeds the runner-up by the requested margin.

**Fig. 4.**
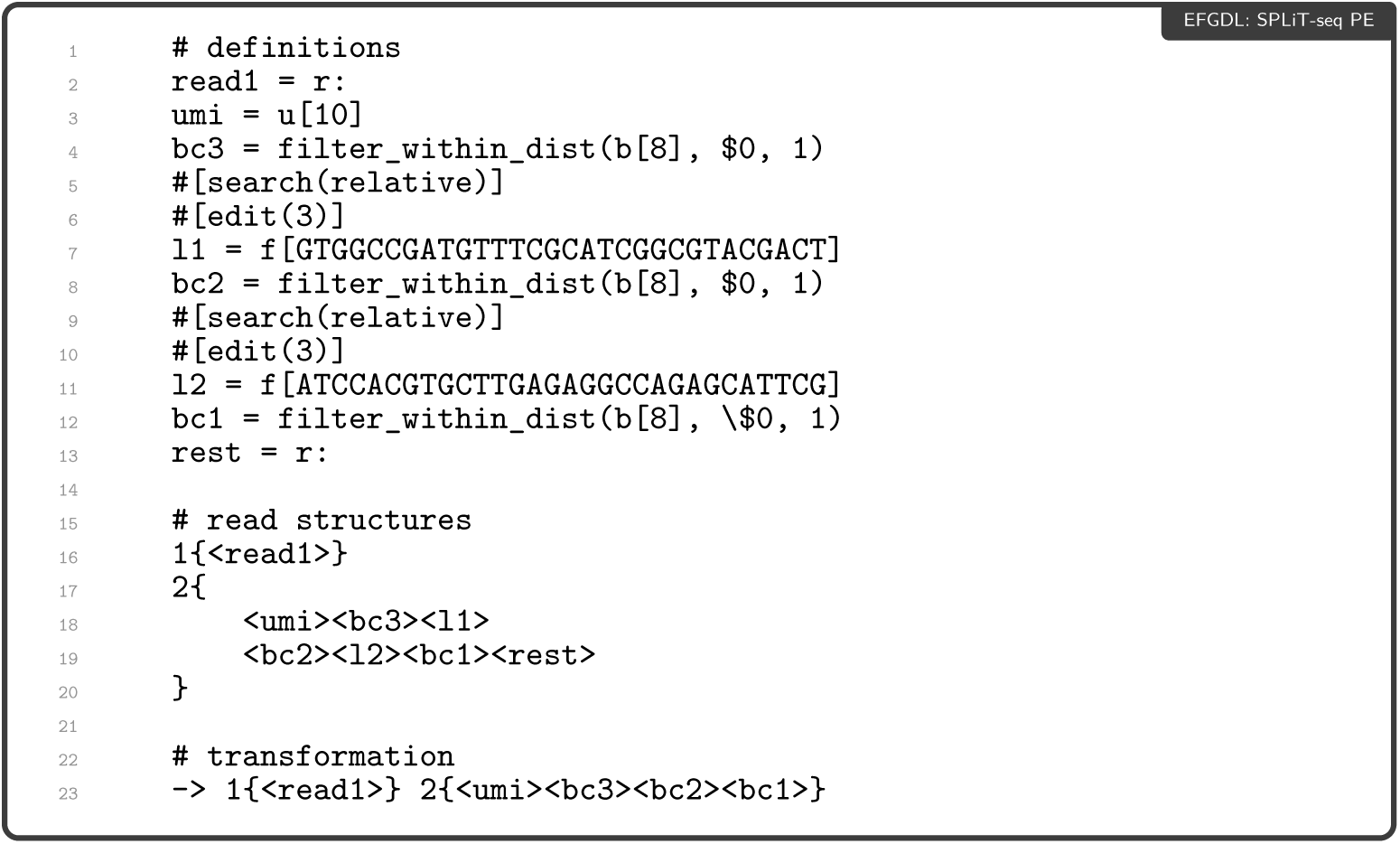
An idiomatic EFGDL specification for the SPLiT-seq PE geometry used in the final benchmark campaign. The input layout is UMI(10)+BC3(8)+Linker 1(30)+BC2(8)+Linker 2(30)+BC1(8), followed by any remaining sequence. Each barcode must be within Hamming distance one of the canonical eight-base list supplied as $0; linkers are located within edit distance three.

The command-line interface separates validation, inspection, and execution. seqproc validate performs semantic validation and reports actionable diagnostics without reading FASTQ data; seqproc explain prints the normalized specification and compiled execution graph; and seqproc run executes the graph. This allows protocol specifications to be checked and reviewed independently of a full processing run.

### 5.3 Descriptive representations of chemistries

To show how definitions, specifiers, annotations, and transformations combine on a real protocol, we walk through a complete EFGDL specification for SPLiT-seq PE data [3], one of the more complex single-cell chemistries. We give the specification in full below and then describe each part.

This EFGDL specification defines the eight intervals used by the paired-end SPLiT-seq geometry. The filter_within_dist transformations require all three eight-base barcodes to occur within Hamming distance one of an entry in the canonical whitelist supplied as command-line argument $0. Linkers l1 and l2 are defined with annotations controlling how they are located. Because SPLiT-seq reads may contain insertions or deletions around the linkers, #[search(relative)] searches for the best fragment match and binds the adjacent intervals relative to it, while #[edit(3)] admits up to three edits, including insertions and deletions. The final transformation retains read 1 unchanged and projects read 2 to the 34-base UMI+BC3+BC2+BC1 product.

### 5.4 ANTISEQUENCE

antisequence is the implementation of the primitives exposed by EFGDL, controlling *how* reads are matched and transformed. It is a sequence recognition and transformation library upon which seqproc is based. We hope that it will be of independent interest beyond its use in seqproc as it has been designed as a general framework for representing matching and transformation of sequencing data.

In antisequence one defines sequences, or more specifically DAGs, of operations that allow recognizing, labeling, and transforming sequencing reads, in addition to associated data such as identifiers and quality scores. These graphs are defined via antisequence’s graph API. Each node in the graph encodes some particular operation to be performed on the incoming sequence, such as labeling a specific segment, trimming or padding a segment, searching the read for a particular pattern (and then labeling it), or even defining some region based on data such as quality scores. Figure 5 illustrates how reads may be operated on in specific antisequence nodes. These nodes can then be connected together via edges that describe the “flow” of reads over the execution graph. In the simplest pipelines, reads flow from one node to the next in a linear fashion, though more complex graphs are also possible wherein the subsequent processing of a read (the path that it takes in the graph) is based on the processing or labeling that occurs in an intermediate node. After being defined, the graph can be optimized prior to execution. For example, certain operations that will always be performed in sequence may be “fused” together to avoid unnecessary overhead. Our current antisequence graph compiler implements some basic optimizations like this, but many more optimization opportunities exist. We believe the links between effective strategies for optimizing execution graphs and optimizing abstract program representations (i.e., as done by different phases of a compiler) are interesting to explore, and we suspect that this general framework could prove very powerful for even complex sequence manipulation tasks.

**Fig. 5.**
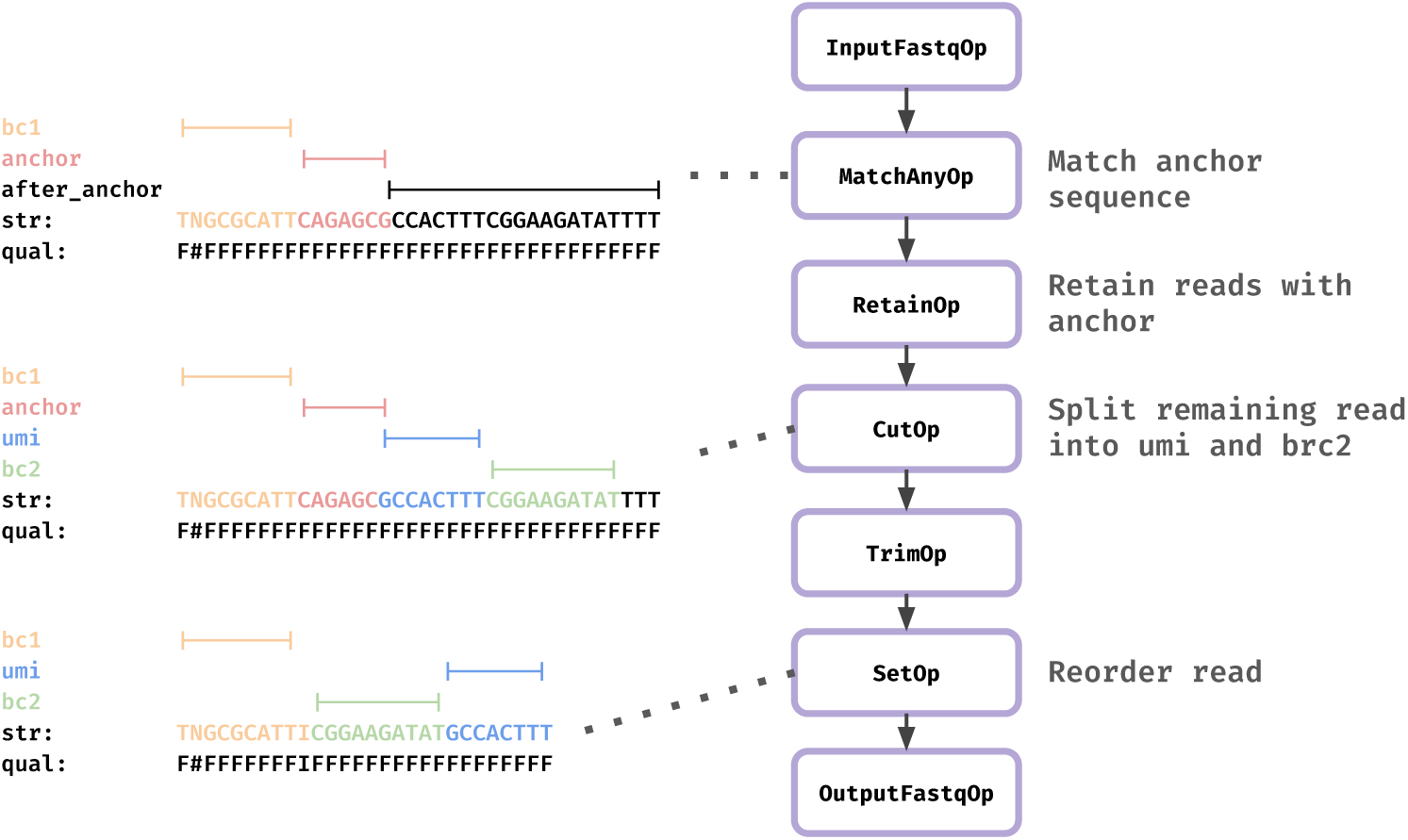
How the EFGDL specification from Figure 3 would be translated into an antisequence graph, as well as how it would match and transform intervals.

After the execution graph is defined and compiled, reads are streamed through it in batches rather than one at a time, with each node operating on a whole batch. Normal worker-local execution uses batches of 512 reads by default; the optional bounded staged pipeline uses 256-read batches by default, and both sizes are configurable. When reads in a batch diverge across a branching graph, keeping the batches coherent raises scheduling questions that trade throughput against working memory. However, as the protocols evaluated here compile to linear graphs, we note this only as a direction for future work (Section 4).

antisequence has been designed with performance as a primary consideration. Approximate matching of short literal patterns (up to 64 bp, which covers barcodes, UMIs, and the linker sequences used by complex chemistries) uses precomputed Myers bit-vector masks [37], whose cost is essentially independent of the edit-distance threshold. Longer literal searches use a multiword Myers implementation, while general dynamically evaluated patterns fall back to dynamic programming. For collections of patterns, a rolling-hash seed index narrows the candidates before verification. Its general path computes eight rotate–XOR rolling hashes per batch and uses the raw HashTable interface from hashbrown, so precomputed 64-bit hashes are not cryptographically re-hashed. Very short, low-distance Hamming matches instead use a precomputed lookup table backed by a fast non-cryptographic hasher. Identity and overlap alignment modes use SIMD (Single Instruction, Multiple Data) acceleration through the block-aligner [38] library, utilizing AVX2 instructions on x86-64 architectures and NEON instructions on ARM (aarch64) platforms. Parallelism uses scoped operating-system threads. The default worker-local mode lets each worker parse and execute a complete graph on its own batches for cache locality. An optional bounded reader–worker–writer pipeline provides backpressure. For workflows that require output reads to appear in the same order as the input, the ––preserve-order flag assigns sequence numbers to batches and restores their order with a bounded reorder buffer while transformations remain parallel. Several mechanisms are used to enhance memory and cache efficiency, including the use of SmallVec for storing mappings, which avoids heap allocations for reads with few labeled intervals (the common case), and the recycling of Read objects between batches to minimize allocation overhead. Additionally, antisequence supports optional high-performance memory allocators (mimalloc and jemalloc) that can provide further throughput improvements depending on workload characteristics.

FASTQ input is parsed with needletail, including transparent gzip decompression. For regular gzip files, an optional rapidgzip-core backend performs adaptive speculative decompression before feeding the same parser and execution graph. Gzip output defaults to compression level 3 and offers two parallel modes: independent batch compression into a standards-compliant concatenated multi-member stream, or background deflate-block compression into one logical gzip member with dictionary continuity. These compressed-I/O options were not used in the uncompressed benchmarks reported here.

### Processing statistics

When enabled via the ––summary flag, seqproc collects runtime statistics and emits a versioned JSON report. A basic report includes *i*) run metadata (version and invocation); *ii*) input, accepted, and rejected read counts; *iii*) rejection reasons and per-input statistics; and *iv*) execution metadata, including ordering mode and effective thread count. Detailed mode additionally records read-length distributions, match-distance histograms for every labeled interval, ordered match-stage attrition, and ambiguity-policy outcomes. The distance histograms track the distribution of distances for accepted matches and the number of unmatched reads, allowing users to evaluate library quality and verify that chosen mismatch thresholds are appropriate. To keep collection overhead low, high-frequency counters such as match distances and read lengths are accumulated per worker and merged only after processing. Match-distance histograms are stored sparsely to support unbounded edit distances without pre-allocating large arrays. Statistics are disabled unless requested, so normal runs do not update these counters. The final summary is serialized to JSON, facilitating downstream analysis and integration with quality control pipelines.

### Abbreviations

ARI, adjusted Rand index; CLI, command-line interface; DAG, directed acyclic graph; DSL, domain-specific language; EFGDL, Extended Fragment Geometry Description Language; LR, long read; PE, paired end; RSS, resident set size; scRNA-seq, single-cell RNA sequencing; UMI, unique molecular identifier.

## Declarations

### Ethics approval and consent to participate

Not applicable.

### Consent for publication

Not applicable.

### Availability of data and materials

The datasets analysed in this study are available from the NCBI Sequence Read Archive under accessions SRR6750041, SRR13948564, SRR8315379, and SRR7827254. The seqproc source code is available at https://github.com/COMBINE-lab/seqproc under the BSD 3-Clause license. The antisequence source code is available at https://github.com/COMBINE-lab/antisequence under the MIT license. Benchmark scripts, configurations, and reproducibility instructions are available at https://github.com/COMBINE-lab/seqproc-paper-analysis.

### Authors’ contributions

**Noah Cape**: Conceptualization; Methodology; Software (lead development of the initial seqproc implementation, including the parser and the seqproc-to-antisequence compiler; EFGDL language and syntax design); Investigation (experimental design and oversight); Writing–original draft; Writing–review and editing.

**Elan Fisher**: Methodology; Software (expansion of seqproc capabilities, including improved matching and scoring backends and the annotation system; optimization of the antisequence backend, including reporting, batching, and execution performance); Investigation (design and execution of the initial experiments); Formal analysis; Validation; Writing–original draft; Writing–review and editing.

**Daniel Liu**: Conceptualization; Methodology; Software (lead development of the initial antisequence library and formulation and implementation of its transformation-graph architecture).

**Rob Patro**: Conceptualization (initial conception of seqproc with Noah Cape); Methodology (design of the DSL-over-antisequence architecture and the syntax and structure of EFGDL); Software (expansion of EFGDL capabilities and further optimization of antisequence); Investigation; Formal analysis (experimental design, implementation, and interpretation); Supervision; Project administration; Writing–original draft; Writing–review and editing.

## Acknowledgements

We thank Dongze He for useful discussions related to the initial design and implementation of seqproc, and Gaurav Sharma for designing the precursor to the FGDL, itself the precursor of the EFGDL.

## Footnotes

1 splitcode also accepts gzip-compressed and interleaved FASTQ input and can emit its output as FASTQ, gzip-compressed FASTQ, unaligned BAM, or interleaved sequences directed to standard output for piping into downstream tools.

2 https://github.com/COMBINE-lab/seqproc-paper-analysis

3 All benchmark scripts, configuration files, and reproducibility instructions are available at https://github.com/COMBINE-lab/seqproc-paper-analysis.

