## Supplementary Information for "SEQPROC: an efficient, flexible, and concise tool for sequence geometry description and transformation"

#### S1 Detecting and handling ambiguity in the EFGDL description

SEQPROC rejects EFGDL specifications that are structurally ambiguous, such as those containing a variable interval left without a right anchor, and resolves the input-dependent ambiguities that even a well-formed specification can admit on a particular read using deterministic tie-breaking rules. For example, the following sequence of intervals describing the first read in a paired-end read specification is ambiguous: `1{b[9-10]r:}2{r:}` and will not be successfully parsed by SEQPROC. For a sequence of length greater than 9, there are two valid ways to associate the variable length barcode and unbounded length read intervals. If, instead, the variable length barcode is anchored on the right by a fragment-specified fixed sequence, for example `1{b[9-10]f[ATGC]r:}2{r:}`, the boundary between the barcode and the read is determined by the position of the fixed sequence. Note that fragment-specified fixed sequence intervals are matched exactly by default. A residual ambiguity can remain when the fixed sequence overlaps itself. For the specification `1{b[3-5]f[ATAT]r:}` and the read `CCGATATATGGGCCC`, the anchor `ATAT` occurs at two offsets compatible with `b[3-5]`, yielding either `b = CCG` or `b = CCGAT`. SEQPROC resolves such cases by taking the leftmost (first) exact occurrence of the fixed sequence, assigning the shortest valid length to the preceding variable interval (here `b = CCG`). Allowing the user to select alternative matching strategies for such cases is a direction for future work (see Section 4).

#### S2 $V_{\text{total}}$ methodology and orthogonal validation

**Definition of $V_{\text{total}}$.** For each protocol, $V_{\text{total}}$ is a protocol-informed, conservative structural reference used to compare the emitted ID set of each method; it is not assumed to contain every biologically recoverable read and is not experimental ground truth. The same strict binary FASTQ parser verifies record structure, paired-file record counts, and mate IDs where applicable. Accepted IDs are streamed in input order rather than accumulated in memory.

| Method | Mean wall time ± SD (s) | | | | Peak RSS at fastest point (MiB) |
|---|---|---|---|---|---|
| | 1 thread | 4 threads | 16 threads | 32 threads | |
| seqproc | **14.650 ± 0.058** | **3.753 ± 0.029** | **2.801 ± 0.029** | **3.118 ± 0.050** | 39.0 |
| matchbox | 420.893 ± 0.703 | 110.146 ± 0.281 | 34.119 ± 0.151 | 21.671 ± 0.307 | **36.0** |
| splitcode | 221.907 ± 1.950 | 57.324 ± 0.603 | 16.554 ± 0.307 | 8.610 ± 0.175 | 532.9 |

Table S1: Controlled forward-orientation-only performance comparison on LR-SPLiT-seq (5,764,421 reads), using the same complete-component and canonical-list constraints as the primary block. The MATCHBOX configuration uses the anchor-first transcription described in the main text. Wall times are means ± sample standard deviations over three randomized runs with uncompressed input and requested output directed to `/dev/null`; peak RSS is the largest replicate maximum resident set size at each method's fastest tested thread count. The capability-complete dual-orientation comparison is the primary LR result in Table 2.

For 10x Chromium v2, a pair enters $V_{\text{total}}$ when both mates are present with matching IDs and R1 has length at least 26; no barcode whitelist is added to this intentionally length-only task. For sci-RNA-seq3, the validator independently evaluates the documented CAGAGC anchor placements after a 9- or 10-base BC1, requires room for UMI(8)+BC2(10), selects a unique minimum-edit placement within edit distance one, and rejects equal-best offset ties. This prevents a better match at an illegal offset from hiding valid protocol structure while making the ambiguity policy explicit.

For paired-end SPLiT-seq, the validator requires complete, ID-matched mate records and the corrected R2 layout UMI(10)+BC3(8) +Linker 1(30)+BC2(8)+Linker 2(30)+BC1(8). Both corrected linkers must match within edit distance three and all three eight-base barcodes must occur within Hamming distance one of their canonical lists. For LR-SPLiT-seq, a read must contain the complete adjacent UMI(10)+BC3(8)+Linker 1+BC2(8)+Linker 2+BC1(6) cassette in either orientation, with linker edit-distance budgets of three and Hamming-distance-one membership in the canonical barcode lists. Both orientations are always evaluated, candidate enumeration is uncapped, and an ID is retained once if either orientation passes. Equal-best barcode owners are retained as structural set membership rather than treated as a unique correction. These choices credit small indels in anchors while requiring a complete, protocol-compatible cassette.

| Chemistry | Accession | $\lvert V_{\text{total}}\rvert$ | % of Total |
|---|---|---|---|
| SPLiT-seq PE | `SRR6750041` | 57,437,503 | 74.00 |
| LR-SPLiT-seq | `SRR13948564` | 560,699 | 9.73 |
| 10x Chromium v2 | `SRR8315379` | 234,382,218 | 100 |
| sci-RNA-seq3 | `SRR7827254` | 19,864,110 | 89.93 |

Table S2: Size of the conservative structural reference ($\lvert V_{\text{total}}\rvert$) for each protocol and its percentage of the complete input. These sets define reference-compatible structure, not biological ground truth.

The smaller LR fraction, 9.73% against 74.00% for PE, reflects the requirement for a complete cassette in the noisier long-read library. It cannot distinguish irrecoverable molecular damage from valid structure excluded by the declared error model. Within the LR core, 1,793 reads pass in both orientations and 129,024 accepted reads have more than one equal-best owner for at least one six-base barcode; these are retained as structurally compatible but are not claimed to have a unique corrected barcode. The uniquely correctable subset contains 431,675 reads. Wider linker-budget sensitivity sets contain 573,518 reads at budgets 5/4 and 590,579 at 6/6, compared with the 560,699-read primary 3/3 core. Reads outside these references remain unadjudicated.

For a tool whose emitted reads form the ID set $E$, the Precision and Recall percentages against $V_{\text{total}}$ are

$$\text{Precision (\%)} = 100 \times \frac{\lvert E \cap V_{\text{total}}\rvert}{\lvert E\rvert},$$

$$\text{Recall (\%)} = 100 \times \frac{\lvert E \cap V_{\text{total}}\rvert}{\lvert V_{\text{total}}\rvert}.$$

and F1 is their harmonic mean; both share the single reference $V_{\text{total}}$ per protocol. The reason-coded Python validator uses ordered multiprocessing, exact-structure fast paths, owner-aware Hamming tables, and streamed ID output, and is run separately from the timed tool comparisons. Figure 2 breaks down, for each dataset, the reads recovered by all three tools, by each pair, and by each tool alone.

#### S3 Downstream concordance details

**Primary canonical-barcode, fuzzy-linker `EditDist_2` analysis.** The analysis of Section 2.3 uses the final, semantically aligned upstream products and one common STARsolo 2.7.11b configuration. The primary matchbox product uses exact canonical-barcode matching, edit-distance-three linkers, and captured-component length guards. We located each tool's barcode-rank cutoff with a Python implementation of the DropletUtils `barcodeRanks` method [1], which fits a smoothing spline to the log–log curve of total UMIs against barcode rank and takes the point of steepest descent as the cell/empty inflection. Applied identically to the three matrices, it places the inflection at rank 258, 255, and 251 for SEQPROC, SPLITCODE, and MATCHBOX (UMI thresholds 101, 109, and 114). Table S3 gives the corresponding per-cell-type label overlap.

| Cell type | SP/SC | SP/MB | SC/MB | Mean |
|---|---|---|---|---|
| Neuron | 0.959 | 0.965 | 0.979 | 0.968 |
| Astrocyte | 0.931 | 0.966 | 0.966 | 0.954 |
| Oligodendrocyte | 0.850 | 1.000 | 0.850 | 0.900 |
| OPC | 0.529 | 0.333 | 0.444 | 0.436 |
| Microglia | 0.625 | 0.444 | 0.529 | 0.533 |
| Endothelial | 0.571 | 0.714 | 0.625 | 0.637 |
| Mean | 0.744 | 0.737 | 0.732 | 0.738 |

Table S3: Pairwise per-cell-type Jaccard index of marker-based labels over the 220 cells called by all three tools. SP, SC, and MB denote SEQPROC, SPLITCODE, and canonical-barcode/fuzzy-linker MATCHBOX. The final column and row retain the across-pair summaries.

**Barcode-correction sensitivity.** The primary `EditDist_2` setting corrects each of the three SPLiT-seq barcode pieces independently. We additionally repeated the complete analysis with the alternative whole-barcode `1MM` correction mode. The upstream read sets are unchanged. For SEQPROC, which preserves the observed barcode bases, the fraction of reads accepted as valid barcodes increases from 99.189% under `1MM` to 99.973% under `EditDist_2`. The corresponding SEQPROC–SPLITCODE per-barcode Pearson correlation increases from 0.972 to 0.981. Called-cell counts remain 225, 220, and 221, while all-tool cell-type agreement changes modestly from 0.891 to 0.909. Thus the correction mode mainly affects recovered counts and some low-signal discrete labels rather than the called-cell population.

**Externally expanded matchbox sensitivity.** The boundary-safe primary MATCHBOX configuration requires exact membership in each 96-member barcode list while permitting fuzzy linker matches under explicit length guards. As a diagnostic upper-bound sensitivity, we externally expanded each list to include every Hamming-distance-one neighbor and repeated both STARSOLO correction modes. Expansion increases the MATCHBOX output from 50,266,141 to 57,252,325 read pairs and its recall against the conservative structural reference from 87.32% to 99.26%; precision changes from 99.78% to 99.58% (Table S4). This is not the primary configuration because generating and managing expanded lists is preprocessing outside MATCHBOX and shifts protocol knowledge from the tool configuration to a generated auxiliary file.

The expanded product calls 225 cells under both `EditDist_2` and `1MM`, compared with 221 for canonical-barcode MATCHBOX. Under `EditDist_2`, expansion raises read-set Jaccard to 0.982 with SEQPROC and 0.922 with SPLITCODE and raises the minimum pairwise per-barcode correlation from 0.962 to 0.975, but it does not qualitatively alter the downstream conclusions (Table S5 and Figure S4). Label-level metrics are not strictly monotone across settings because marker-based typing and independent Leiden clustering discretize relatively small expression differences, particularly for rare cell types. We therefore treat the consistently high gene-level correlations and stable major cell populations as the more robust result and present the expanded configuration only as a capability sensitivity.

| MATCHBOX barcode list | Emitted reads | Precision (%) | Recall (%) | F1 |
|---|---|---|---|---|
| Canonical barcodes | 50,266,141 | 99.78 | 87.32 | 0.9313 |
| Hamming-1-expanded barcodes | 57,252,325 | 99.58 | 99.26 | 0.9942 |

Table S4: Structural-reference sensitivity for canonical and externally Hamming-1-expanded MATCHBOX barcode lists, both with edit-distance-three linker matching and captured-component length guards. Precision and recall are agreement with the conservative structurally retained set, not biological ground truth.

**Quality-score interpretation.** The three upstream programs do not write equivalent barcode qualities. SEQPROC preserves observed barcode bases and their measured quality scores. The SPLITCODE replacement workflow writes canonical barcode bases and synthetic (fixed) qualities for those barcode segments, while preserving the UMI quality. Canonical-barcode MATCHBOX emits observed bases that are canonical by construction and retains observed qualities; its expanded-list sensitivity also retains the observed one-mismatch variants and their qualities. Consequently, the STARSOLO Q30 summary for CB+UMI is not a comparable accuracy metric across tools. The present `1MM` and `EditDist_2` comparisons quantify the effect of correction geometry and upstream retention. They should not be interpreted as a controlled comparison of quality-aware correction. Preserving measured qualities remains a useful native capability of SEQPROC and MATCHBOX when a downstream method can use them.

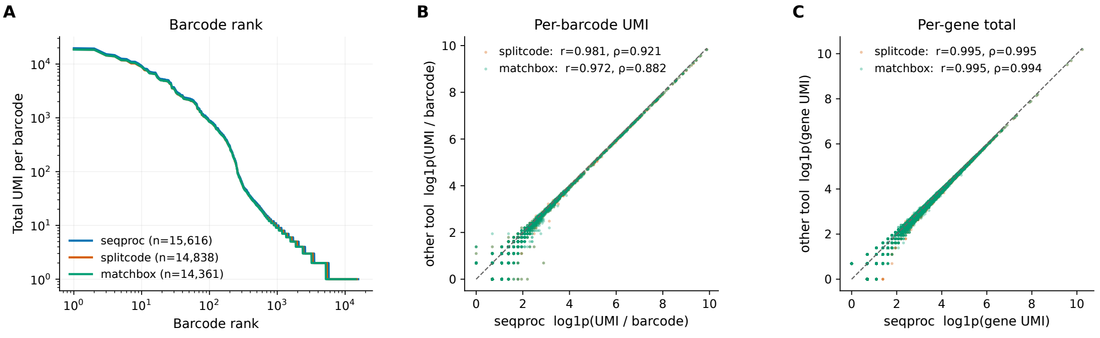


Figure S1: Count concordance for the primary canonical-barcode, fuzzy-linker `EditDist_2` analysis. **(A)** Barcode-rank curves contain 15,616, 14,838, and 14,361 nonzero barcodes, with inflections at ranks 258, 255, and 251 for SEQPROC, SPLITCODE, and MATCHBOX. **(B)** Per-barcode total UMI on the $\log(1+x)$ scale. Pearson correlation is 0.981 for SEQPROC–SPLITCODE and 0.972 for SEQPROC–MATCHBOX (Spearman 0.921 and 0.882). **(C)** Per-gene totals are more concordant: Pearson correlation is 0.995 for both pairs and Spearman correlation is 0.995 and 0.994, respectively.

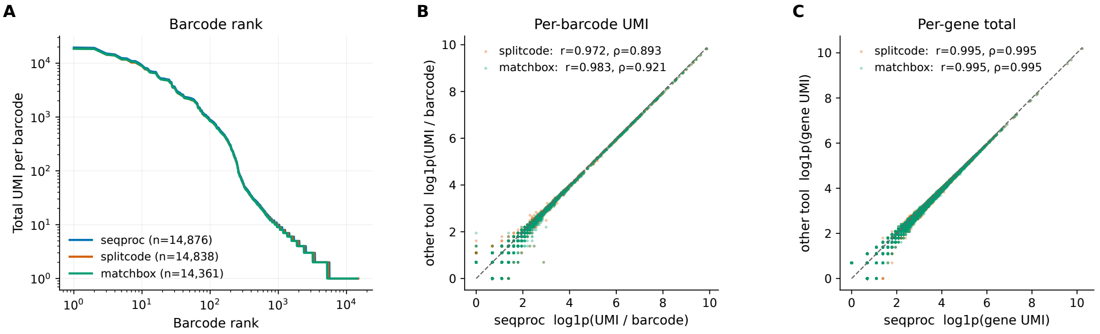


Figure S2: Canonical-barcode `1MM` sensitivity analysis. **(A)** The nonzero matrices contain 14,876, 14,838, and 14,361 barcodes, with inflections at ranks 257, 255, and 251. **(B)** Per-barcode Pearson correlation is 0.972 for SEQPROC–SPLITCODE and 0.983 for SEQPROC–MATCHBOX. **(C)** Per-gene Pearson correlation is 0.995 for both pairs.

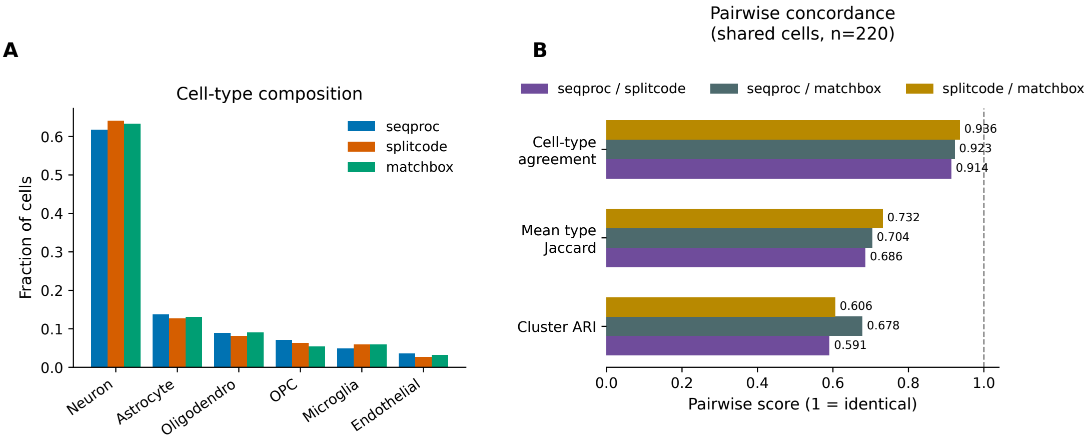


Figure S3: Biological concordance for the canonical-barcode `1MM` sensitivity. **(A)** Cell-type composition among the 225, 220, and 221 cells called for SEQPROC, SPLITCODE, and MATCHBOX. **(B)** Pairwise concordance on the 220 shared cells. For SEQPROC–SPLITCODE, SEQPROC–MATCHBOX, and SPLITCODE–MATCHBOX, respectively, cell-type agreement is 0.914, 0.923, and 0.936; mean per-type Jaccard is 0.686, 0.704, and 0.732; and cluster ARI is 0.591, 0.678, and 0.606. The complementary all-tool cell-type agreement is 0.891 and joint co-clustering agreement is 1.000.

| Configuration | Matchbox reads | Valid BC (%) | Cells (SP/SC/MB) | All-type | Mean Jaccard | Min. gene $r$ | Min. barcode $r$ |
|---|---|---|---|---|---|---|---|
| Canonical + `EditDist_2` | 50,266,141 | 99.97 | 225/220/221 | 0.909 | 0.738 | 0.991 | 0.962 |
| Canonical + `1MM` | 50,266,141 | 99.97 | 225/220/221 | 0.891 | 0.708 | 0.991 | 0.962 |
| Expanded + `EditDist_2` | 57,252,325 | 99.97 | 225/220/225 | 0.900 | 0.747 | 0.995 | 0.975 |
| Expanded + `1MM` | 57,252,325 | 99.18 | 225/220/225 | 0.891 | 0.714 | 0.995 | 0.968 |

Table S5: Sensitivity of the downstream analysis to STARsolo barcode correction and to external Hamming-1 expansion of the canonical MATCHBOX barcode lists. All MATCHBOX variants use edit-distance-three linker matching and captured-component length guards. SP, SC, and MB denote SEQPROC, SPLITCODE, and MATCHBOX. "Min." is the least favorable value among the three tool pairs, avoiding post hoc selection of a favorable comparison.

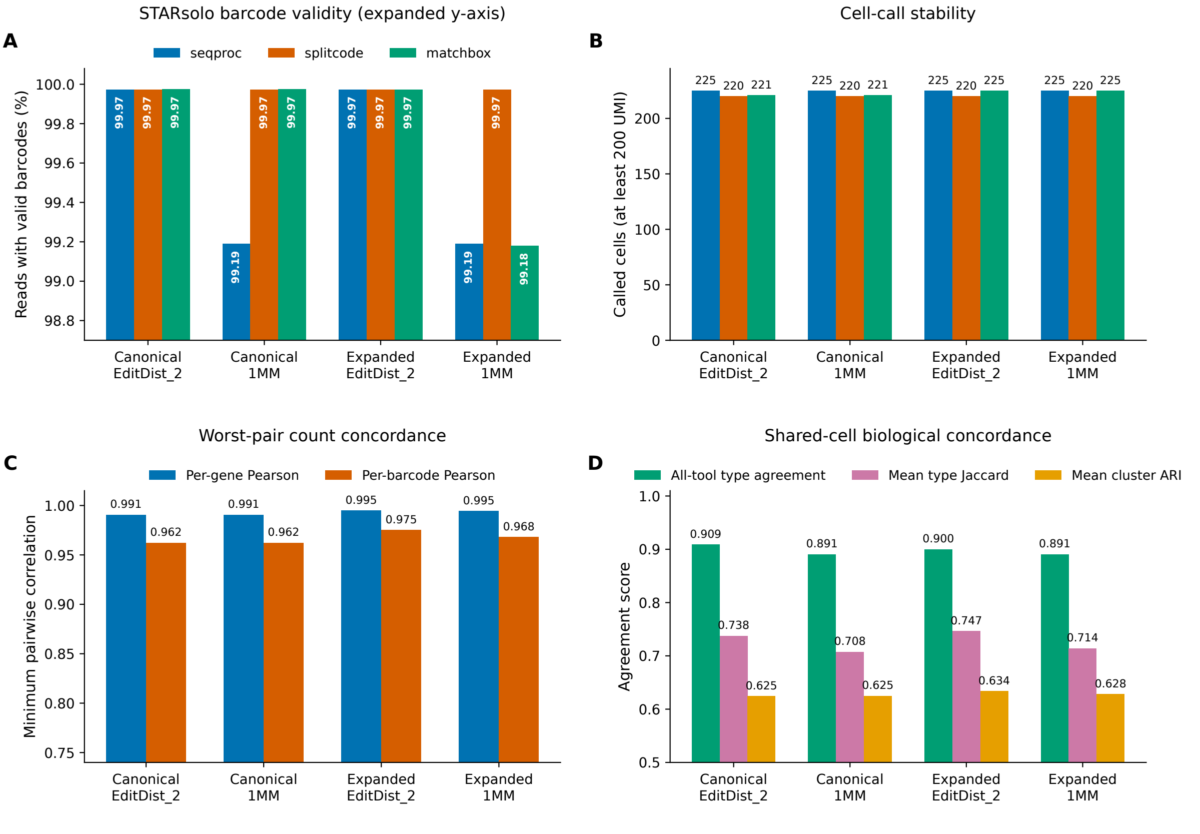


Figure S4: Downstream sensitivity across canonical and externally Hamming-1-expanded MATCHBOX lists and the two STARsolo correction modes. All MATCHBOX configurations use fuzzy linkers and captured-component length guards. **(A)** Reads with valid barcodes; the y-axis is expanded to expose differences near 100%. **(B)** Cells called with at least 200 UMIs. **(C)** The minimum pairwise Pearson correlation among all three tool pairs for per-gene and per-barcode totals. **(D)** Compact across-pair summaries on shared called cells: all-tool cell-type agreement, mean per-type Jaccard, and mean pairwise cluster ARI. The constituent pairwise values are reported in Table S6.

| Configuration | Pair | Cell-type agreement | Mean type Jaccard | Cluster ARI |
|---|---|---|---|---|
| | SP/SC | 0.936 | 0.744 | 0.634 |
| Canonical + `EditDist_2` | SP/MB | 0.932 | 0.737 | 0.635 |
| | SC/MB | 0.936 | 0.732 | 0.606 |
| | SP/SC | 0.914 | 0.686 | 0.591 |
| Canonical + `1MM` | SP/MB | 0.923 | 0.704 | 0.678 |
| | SC/MB | 0.936 | 0.732 | 0.606 |
| | SP/SC | 0.936 | 0.744 | 0.634 |
| Expanded + `EditDist_2` | SP/MB | 0.936 | 0.766 | 0.724 |
| | SC/MB | 0.923 | 0.731 | 0.545 |
| | SP/SC | 0.914 | 0.686 | 0.591 |
| Expanded + `1MM` | SP/MB | 0.936 | 0.724 | 0.685 |
| | SC/MB | 0.927 | 0.732 | 0.608 |

Table S6: Pairwise shared-cell biological concordance underlying the aggregate sensitivity scorecard in Figure S4. SP, SC, and MB denote SEQPROC, SPLITCODE, and MATCHBOX. Mean type Jaccard is the unweighted mean over the six marker-based cell types.

### S4 EFGDL transformations and annotations

Table S7 outlines the native transformations and annotations available for specifying protocols in EFGDL. Most can be composed together to keep a specification succinct while allowing one to create complicated transformations.

| **Function** | **Arguments** | **Description** |
|---|---|---|
| `rev(I)` | `I`: interval | Reverse the matched interval. |
| `revcomp(I)` | `I`: interval | Replace the matched interval with its reverse complement (non-ATGC characters unchanged). |
| `trunc(I, n)` | `I`: interval; `n`: positive integer | Truncate the interval by `n` characters from the right. |
| `trunc_left(I, n)` | `I`: interval; `n`: positive integer | Truncate the interval by `n` characters from the left. |
| `trunc_to(I, n)` | `I`: interval; `n`: positive integer | Truncate the interval to length `n` from the right. |
| `trunc_to_left(I, n)` | `I`: interval; `n`: positive integer | Truncate the interval to length `n` from the left. |
| `remove(I)` | `I`: interval | Remove the matched interval. |
| `pad(I, n, c)` | `I`: interval; `n`: positive integer; `c`: character | Pad the interval with `n` copies of `c` on the right. |
| `pad_left(I, n, c)` | `I`: interval; `n`: positive integer; `c`: character | Pad the interval with `n` copies of `c` on the left. |
| `pad_to(I, n, c)` | `I`: interval; `n`: positive integer; `c`: character | Pad the interval to length `n` with character `c` on the right. |
| `pad_to_left(I, n, c)` | `I`: interval; `n`: positive integer; `c`: character | Pad the interval to length `n` with character `c` on the left. |
| `norm(I)` | `I`: variable length interval | Normalize a variable length interval with lower bound (lb) and upper bound (ub) on length to a fixed length interval of length $\mathrm{ub} + \lceil(\lfloor\log_2(\mathrm{ub} - \mathrm{lb} + 1)\rfloor + 1)/2\rceil$, where the appended symbols encode the original length. |
| `map(I, m, f)` | `I`: interval; `m`: mapping file; `f`: fallback expression | Map intervals to sequences in the mapping file; evaluate the fallback expression for unmapped intervals. |
| `map_with_mismatch(I, m, f, n)` | `I`: interval; `m`: mapping file; `f`: fallback expression; `n`: Hamming distance | Map intervals allowing Hamming distance `n`; evaluate the fallback expression for unmapped intervals. |
| `map_with_edit(I, m, f, n)` | `I`: interval; `m`: mapping file; `f`: fallback expression; `n`: edit distance | Map intervals allowing edit distance `n`, tolerating insertions and deletions; evaluate the fallback expression for unmapped intervals. |
| `filter(I, f)` | `I`: interval; `f`: sequence file | Filter the interval against the file, keeping only reads that pass. |
| `filter_within_dist(I, f, n)` | `I`: interval; `f`: sequence file; `n`: positive integer | Filter the interval allowing Hamming distance `n`, keeping only reads that pass. |
| `#[hamming(n)]` | `n`: positive integer | Match the fragment interval with Hamming distance `n`. Applied as an annotation on a definition. |
| `#[edit(n)]` | `n`: positive integer | Match the fragment interval with edit distance `n`, tolerating insertions and deletions. Applied as an annotation on a definition. |
| `#[search(relative)]` | | Search for an anchor sequence anywhere in the read and extract preceding elements with flexible length relative to the anchor position. Can be combined with `#[edit(n)]` or `#[hamming(n)]`. |
| `#[match_ori(either)]` | | Match the read in either orientation, trying the forward orientation first and reverse-complementing the read on failure. Applied as an annotation on a read, important for protocols with random read orientation such as LR-SPLiT-seq. |
| `#[ambig_policy = p]` | `p`: `accept`, `first`, `no_match`, `random(seed = N)`, `quality(min_delta = N)`, or `error` | Resolve an equal-best match against multiple distinct whitelist or mapping entries. Applied to the definition containing the filter or mapping operation. |

Table S7: EFGDL transformations and annotations. Matching modifiers (`#[hamming]`, `#[edit]`, `#[search]`) and ambiguity policy are specified as annotations on interval definitions, and `#[match_ori]` is specified on a read. Other transformations use function syntax applied to intervals within read structures.
